# Role of microRNA-145 in DNA damage signalling and senescence in vascular smooth muscle cells of Type 2 diabetic patients

**DOI:** 10.1101/2020.08.12.248542

**Authors:** Karen E Hemmings, Kirsten Riches-Suman, Marc A Bailey, David J O’Regan, Neil A Turner, Karen E Porter

## Abstract

Increased cardiovascular morbidity and mortality in individuals with type 2 diabetes (T2DM) is a significant clinical problem. Despite advancements in achieving good glycaemic control, this patient population remains susceptible to macrovascular complications. We previously discovered that vascular smooth muscle cells (SMC) cultured from T2DM patients exhibit persistent phenotypic aberrancies distinct from those of individuals without a diagnosis of T2DM. Notably, persistently elevated expression levels of microRNA-145 co-exist with characteristics consistent with aging, DNA damage and senescence. We hypothesised that increased expression of microRNA-145 plays a functional role in DNA damage signalling and subsequent cellular senescence specifically in SMC cultured from the vasculature of T2DM patients. In this study, markers of DNA damage and senescence were unambiguously and permanently elevated in native T2DM versus non-diabetic (ND)-SMC. Exposure of ND cells to the DNA-damaging agent etoposide inflicted a senescent phenotype, increased expression of apical kinases of the DNA damage pathway and elevated expression levels of microRNA-145. Overexpression of microRNA-145 in ND-SMC revealed evidence of functional links between them; notably increased secretion of senescence-associated cytokines and chronic activation of stress-activated intracellular signalling pathways, particularly the mitogen-activated protein kinase, p38α. Exposure to conditioned media from microRNA-145 overexpressing cells resulted in chronic p38α signalling in naïve cells, evidencing a paracrine induction and reinforcement of cell senescence. We conclude that targeting of microRNA-145 may provide a route to novel interventions to eliminate DNA-damaged and senescent cells in the vasculature and to this end further detailed studies are warranted.

## Introduction

The mechanisms underlying increased risk of coronary heart disease (CHD) and its attendant complications in Type 2 diabetes (T2DM) are poorly understood. T2DM reportedly confers up to 15 years of aging (1) whilst chronological aging, characterised by loss of replication-competent cells and DNA damage, is a major risk factor for CHD. It is well known that overwhelming DNA damage triggers a permanent DNA damage response (DDR) and consequent cellular senescence, a permanent arrest of cell proliferation that is a fundamental mechanism of aging that may also play a role in some of the cardiovascular complications of T2DM (2).

Patients with significant CHD frequently require arterial reconstruction in the form of coronary artery bypass grafting (CABG), which in individuals with diabetes is most frequently performed using autologous saphenous vein (SV) grafts (3,4). Phenotypic modulation of the venous smooth muscle cells (SMC) is an important mechanism that enables vein graft adaptation to an arterial environment. Indeed, effective integration to an environment of increased flow and pressure early after implantation is a key determinant of the long-term patency of SV grafts (5). SMC are not terminally differentiated and it is their ability to retain plasticity, i.e. to modulate their phenotype in response to environmental cues, which is key to their role in maintaining vascular homeostasis. Our group previously discovered a distinct and persistent aberrant phenotype, specifically in SV-SMC cultured from T2DM patients compared with those cultured from age-matched non-diabetic (ND) patients (6-8). The enlarged, flattened morphology and impaired proliferation of T2DM-SMC bore resemblance to those of aged, DNA damaged and senescent cells. However, the deleterious effects of DNA damage, senescence and subsequent acquisition of a senescence-associated secretory phenotype (SASP) in SV-SMC, specifically in the setting of human T2DM, have not been explored.

More recently we pursued the notion that persistence of a T2DM SV-SMC phenotype in long-term culture was suggestive of “metabolic memory” underpinned by an epigenetic mechanism. Indeed, we demonstrated that this phenotype was driven by elevated expression levels of a specific SMC-enriched microRNA, miRNA-145 (8). Some recent studies have shown that miRNAs, short non-coding RNAs that are negative regulators of gene expression, in addition to regulating a host of cellular functions, also play crucial roles in cellular responses to DNA damage (reviewed in (9)). Dysregulated miRNAs are also believed to contribute to accelerated aging syndromes (10) and some vascular pathologies (11-13), yet little is known about any relationship between miRNAs, diabetes and vascular SMC dysfunction. Lack of consistency among cell types has suggested that dysregulation of miRNAs after DNA damage is cell-type specific (14). The persistent aberrant phenotype and accompanying elevated levels of miRNA-145 that we regularly observe in T2DM SV-SMC (8) led us to hypothesise that a miRNA-145-mediated mechanism underlies defective DNA damage/repair pathways and subsequent cellular senescence, specifically in this cell-type.

## Results

### Inherent characteristics of senescence and DNA damage in native SV-SMC

We previously reported impaired proliferation and an enlarged flattened morphology characteristic of senescence, in SV-SMC cultured from T2DM patients (6) which persists in culture and throughout passaging (8). Given that there is no single and definitive hallmark of senescence (15), we examined a number of additional endpoints. T2DM-SMC exhibited significantly increased senescence-associated (SA) β-gal staining (Figure 1A), increased expression of IL-1α mRNA (Figure 1B), and reduced expression of nuclear lamin B1 mRNA (LMNB1) (16) (Figure 1C), relative to ND-SMC. No differences were observed in apoptosis (caspase-3 fluorescence assay) (Figure 1D). In addition, a significantly increased frequency of γH2AX positive nuclei, symbolic of double-stranded DNA breaks (Figure 1E) and aberrant nuclear morphology (Figure 1F) in the T2DM cells was indicative of augmented DNA damage. Expression of IL-8 mRNA, an inflammatory mediator associated with SASP, was elevated; however additional SASP mediators IL-6 and MCP-1 were unchanged at the mRNA level (Figure 1G-I). Secretion of IL-6, IL-8 and MCP-1 from native ND and T2DM-SMC was not significantly different but considerable variability between patients was observed, independent of diabetic status (Supplementary Figure 1). These data provide evidence that SV-SMC cultured from patients with T2DM exhibit inherently higher levels of senescence and DNA damage than those of ND patients.

**Figure 1.**
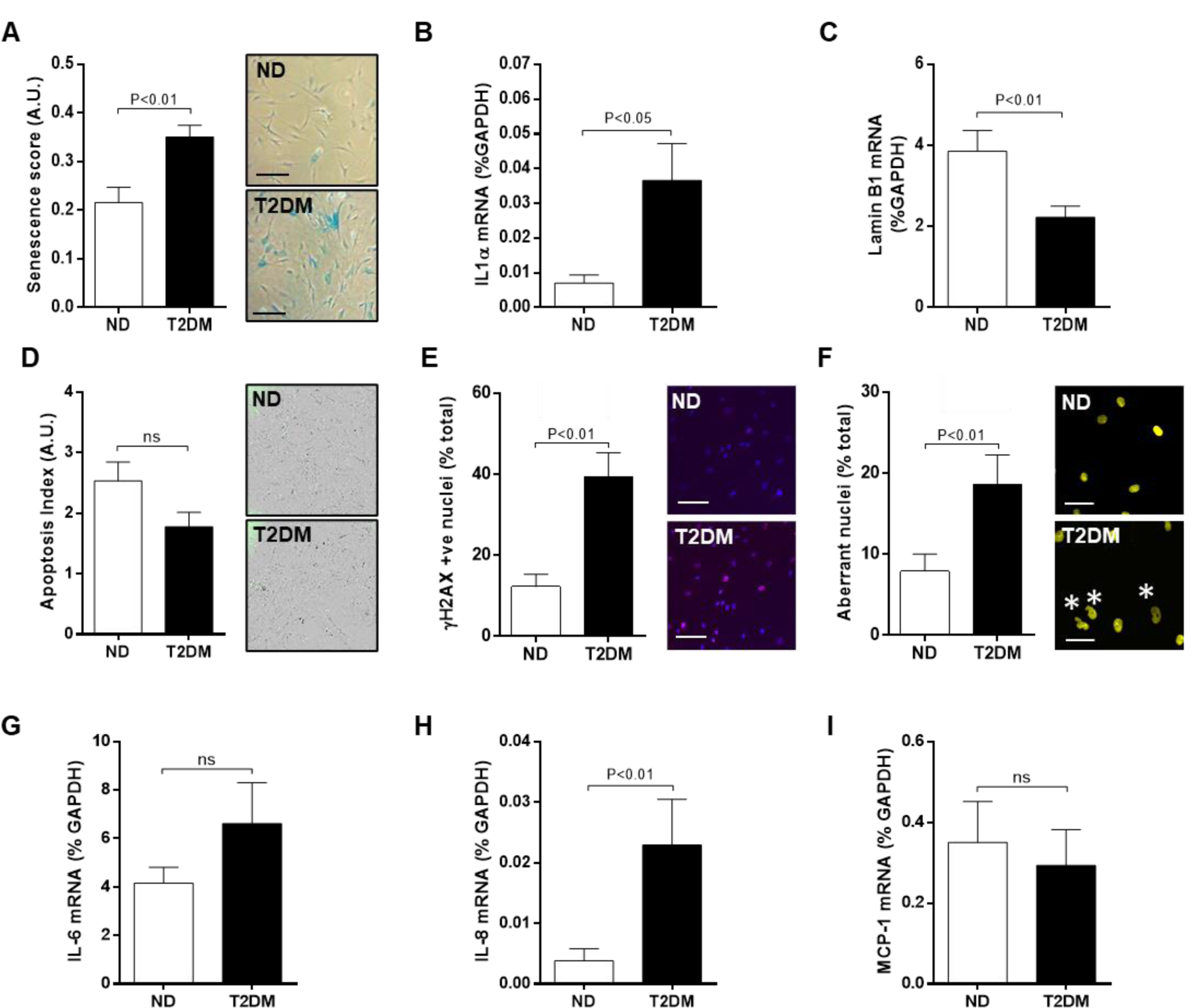
T2DM-SMC exhibit senescent features. **A.** SMC were cultured in full growth medium for 48 h, fixed and stained for senescence-associated β-galactosidase (blue precipitate, scale bar = 200 μm, n=7). **B**. RNA was extracted from cells cultured for 24 h and expression of senescence-associated markers IL-1α and **C**. LMNB1 measured by RT-PCR (n=8). **D**. SMC were cultured in full growth medium for 48 h, fixed and incubated with NucView 488 caspase 3 substrate (to indicate apoptotic cells, n=4), **E.** stained for early DNA damage marker γH2AX (pink foci, scale bar = 100 μm, n=8), and **F.** DAPI to visualise aberrant nuclei (denoted by asterisk, scale bar = 50 μm, n=8). Basal expression of senescence-associated inflammatory genes (G) IL-6, (H) IL-8 and (I) MCP-1 was measured using RT-PCR (all n=8).

### DNA damage and DDR pathway activation in SV-SMC

We next examined whether DNA damage alone could drive the acquisition of a T2DM-SMC phenotype in ND-SMC. Exposure to environmental stresses induces activation of the DDR signalling pathway, a cascade of kinase activations which culminate in cell cycle arrest to enable DNA repair. Phosphorylation of apical kinases ATM (ataxia-telangiectasia mutated), ATR (ATM- and Rad3-Related) and/or DNA-PK (DNA-dependent protein kinase catalytic subunit), depending on the type of DNA damage, activates multiple downstream events that lead to phosphorylation of H2AX (γH2AX) and the tumour suppressor gene p53, and expression of the cell cycle inhibitor p21 that confers growth arrest (Figure 2A).

**Figure 2.**
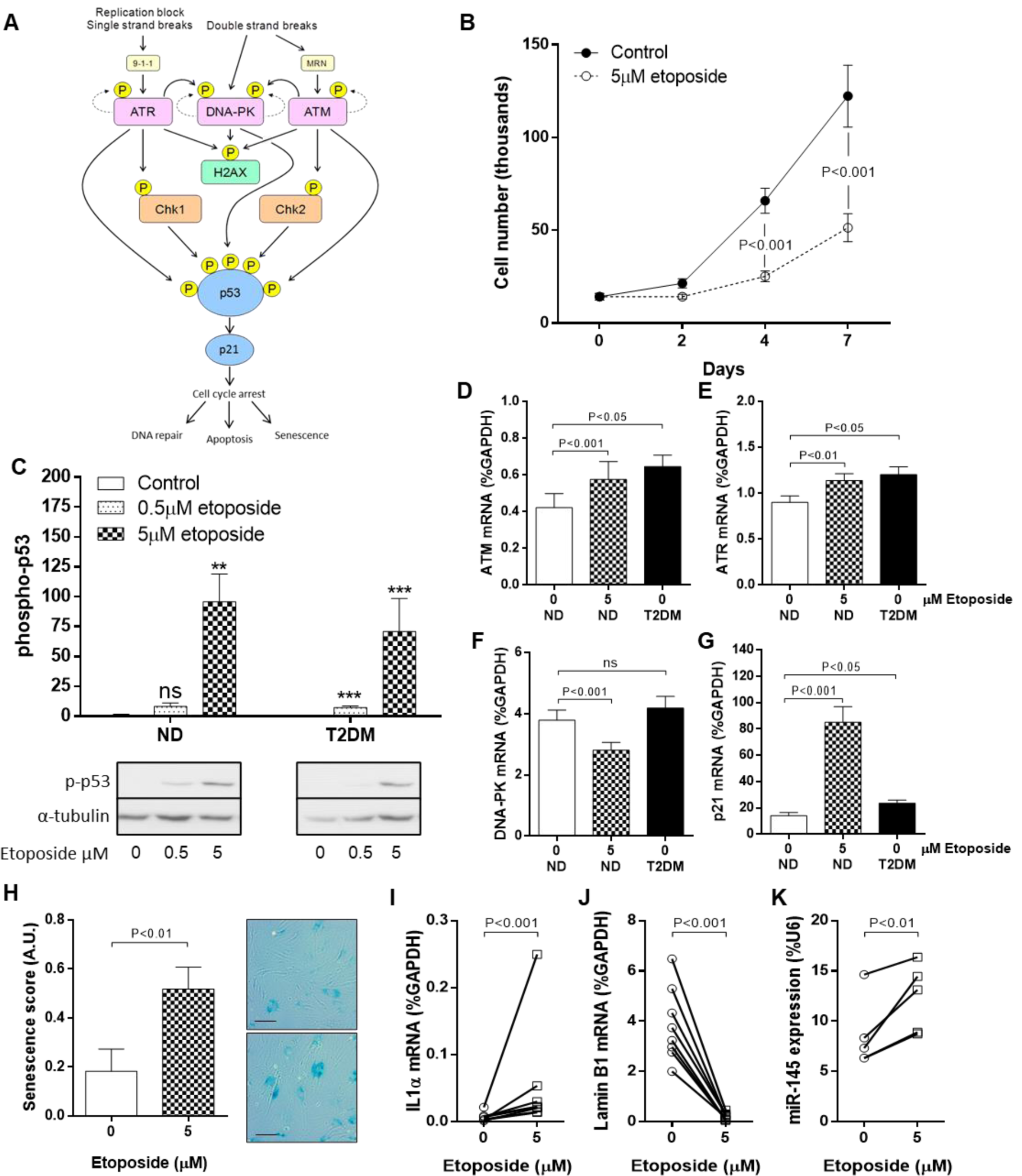
DNA damage drives a senescent SMC phenotype. **A.** The DNA damage signalling pathway leading to phosphorylation of p53, and subsequent downstream effects. **B.** SMC were treated with DNA-damaging agent etoposide (5 μM) or vehicle control (DMSO) for up to 7 days and proliferation quantified by cell counting (n=7). **C.** The impact of etoposide on p53 phosphorylation was monitored using Western blotting after 24 h (n=4). **D.** ND-SMC were treated with 5 μM etoposide for 24 h and the expression of ATM, **E.** ATR, **F.** DNA-PK and **G.** p21 in comparison to vehicle-treated T2DM-SMC measured using RT-PCR (all n=8). **H.** Cells were treated with etoposide for 24 h and then placed into FGM for 72 h. Cells were stained with SA-β-galactosidase (scale bar = 100μm). **I.** The influence of etoposide on expression of IL-1α, **J.** LMNB1 (both n=8) and **K.** miRNA-145 expression (n=6-8) was quantified using RT-PCR.

DNA damage was induced in ND-SMC using the chemotherapeutic agent etoposide (Sigma-Aldrich), which led to a marked reduction in cell proliferation (Figure 2B) and increased phosphorylation of p53 (Figure 2C). Whilst ATM and ATR mRNA were expressed basally at higher levels in T2DM-SMC than in ND cells, exposure of ND cells to etoposide induced increases in both ATM and ATR mRNA, comparable to the levels of expression in native T2DM-SMC (Figure 2D,E). Conversely, DNA-PK mRNA did not differ between ND and T2DM-SMC and etoposide treatment caused a reduction in DNA-PK mRNA expression (Figure 2F). Basal gene expression of p21 was inherently higher in T2DM cells and etoposide treatment of ND cells led to a marked increase in p21 mRNA levels (Figure 2G). Markers of senescence, namely increased SA β-gal staining (Figure 2H), increased IL-1α mRNA expression (Figure 2I) and reduced LMNB1 mRNA expression (Figure 2J), were observed in ND-SMC in response to etoposide, consistent with features detected in native T2DM-SMC (Figure 1A-C). Finally, ND-SMC treated with etoposide exhibited a 1.5-fold increase in miRNA-145 expression (Figure 2K), consistent with differences we previously reported between native ND- and T2DM-SMC in a cohort of 130 patients (8). miRNA-145 expression was also induced by oxidative stress, a more physiologically relevant stimulus (Supplementary Figure 2).

### Pharmacological inhibition of DDR kinases

To examine whether elevated DDR expression was responsible for the T2DM-SMC phenotype, we employed a pharmacological approach in ND-SMC to investigate the effect of apical kinase inhibition on SMC proliferation. Whilst inhibition of ATM (KU55933) or DNA-PK (NU7026) did not modulate proliferation (Figure 3A,C), ATR inhibition (AZ20) completely abolished any increase in cell number over a 7-day period (Figure 3B). miRNA-145 expression was unaffected by any of the inhibitors (Figure 3D). Surprisingly, inhibition of ATM or ATR *increased* ATM mRNA expression (Figure 3E). None of the three inhibitors affected ATR mRNA expression levels (Figure 3F) and inhibition of ATM significantly increased DNA-PK gene expression by 23% (Figure 3G). Inhibition of ATR suggested a possible trend to increased p21 mRNA expression (Figure 3H).

**Figure 3.**
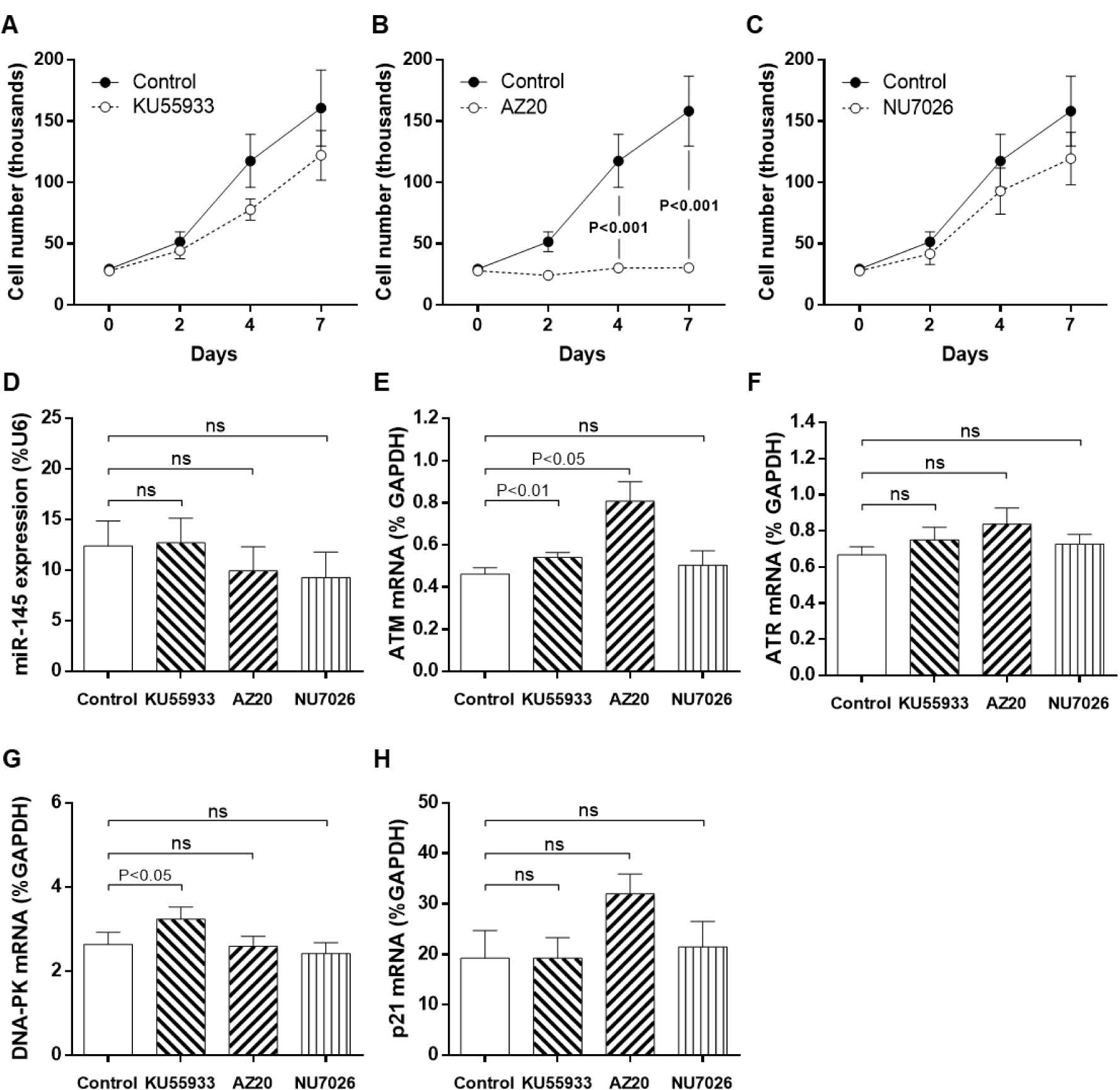
Pharmacological inhibition of apical kinases of the DNA damage response pathway modulates proliferation and gene expression. **A.** SMC were treated with ATM inhibitor (KU55933; 10μM), **B.** ATR inhibitor (AZ20; 10μM) or **C.** DNA-PK inhibitor (NU7026; 3μM) for up to 7 days and cell counts performed on days 0, 2, 4 and 7 (n=4). After 72 h, RNA was extracted and the effect of all inhibitors on expression of **D.** miRNA-145, **E.** ATM, **F.** ATR, **G.** DNA-PK and **H.** p21 was measured using RT-PCR (all n=6). ns = non-significant.

### siRNA knockdown of DDR kinases

To confirm these pharmacological data, we employed a gene-silencing approach to individually knock down ATM, ATR and DNA-PK. siRNA inhibition caused selective and specific reduction of each target mRNA within 24 h and was maintained for at least 96 h (Supplementary Figure 3). Anti-proliferative effects were comparable with pharmacological inhibition, specifically that ATM and DNA-PK knockdown did not modulate SMC proliferation (Figure 4A,C) whilst ATR knockdown led to a significant reduction in cell number (Figure 4B). As with pharmacological inhibition, miRNA-145 expression was unaffected by ATM, ATR or DNA-PK knockdown (Figure 4D). Notably, the gene silencing approach enabled selective inhibition of each kinase, without off-target effects on the others (Figure 4E-G). In agreement with the pharmacological inhibition data, the effect of ATR knockdown on p21 expression was again negligible, although increased p21 mRNA expression was observed as a result of DNA-PK silencing (Figure 4H).

**Figure 4.**
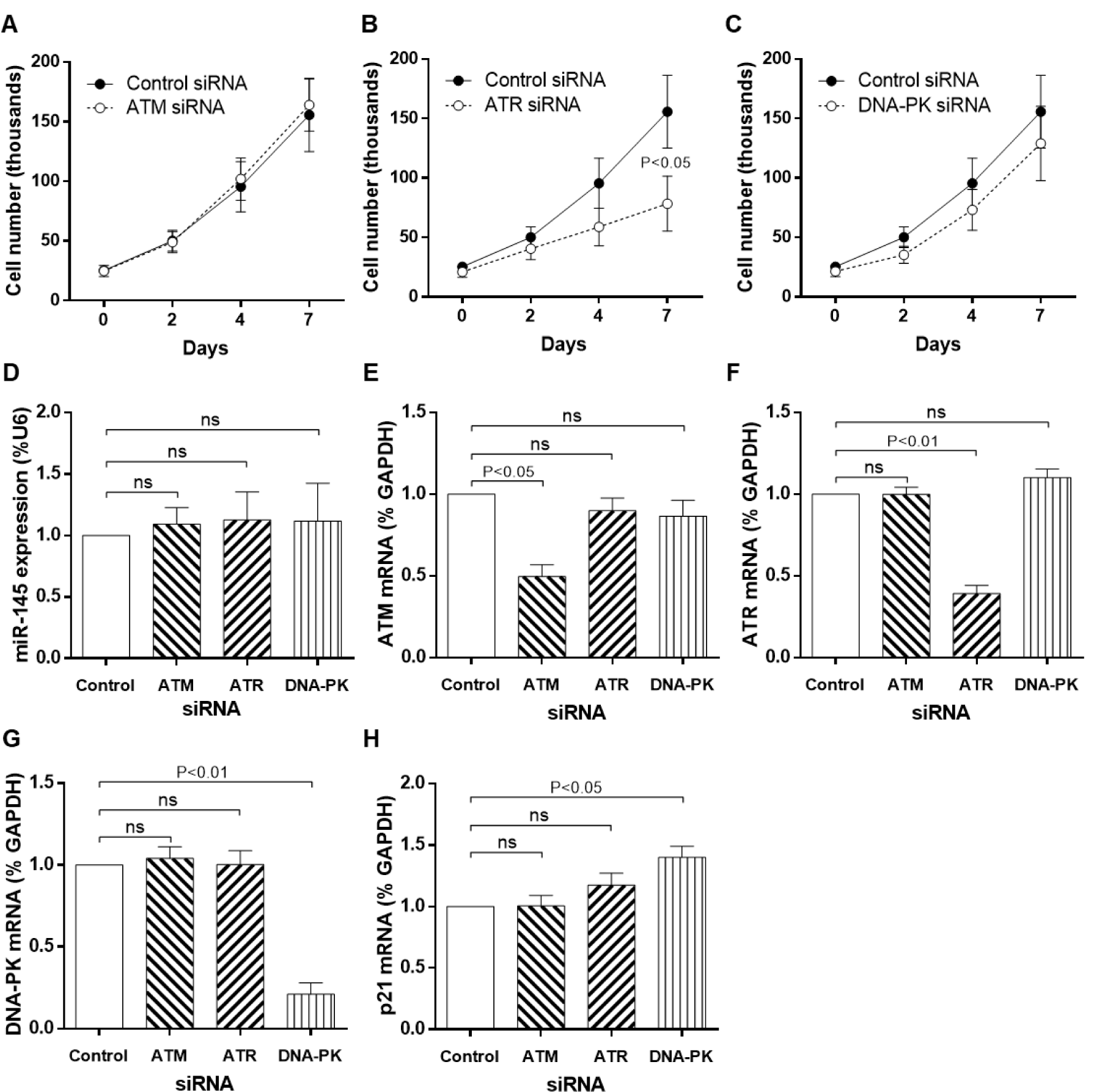
siRNA knockdown of apical kinases of the DNA damage response pathway modulates proliferation and gene expression. **A.** SMC were transfected with siRNA targeted to ATM, **B.** ATR or **C.** DNA-PK for 6 h and then placed into FGM for up to 7 days. Cell counts were performed on days 0, 2, 4 and 7 (n=4). RNA was extracted at 72 h and the expression of **D.** miRNA-145, **E.** ATM, **F.** ATR, **G.** DNA-PK and **H.** p21 measured using RT-PCR (n=5).

### miRNA-145 overexpression modulates DNA damage signalling and senescence

To determine whether miRNA-145 was upstream or downstream of DDR, ND-SMC were transfected with premiRNA –ve (negative control) or premiRNA-145 (overexpression) and parameters pertinent to the DNA damage pathway were evaluated. Overexpression of miRNA-145 led to increased SA β-gal 96 h post transfection (Figure 5A), although differences in γH2AX foci were less evident (P=0.08, Figure 5B). In contrast to increased IL-1α and reduced LMNB1 gene expression observed in native T2DM cells, the opposite was observed in miRNA-145 overexpressing cells; *reduced* IL-1α mRNA and *increased* LMNB1 mRNA (Figure 5C,D). With respect to apical kinases, whilst no consistent differences in ATM gene expression were observed (Figure 5E), increased mRNA expression of ATR (Figure 5F) and decreased mRNA expression of DNA-PK (Figure 5G) were clear. Changes in p21 expression following miRNA-145 transfection were inconsistent (Figure 5H). Importantly, secreted levels of IL-6 (52% increase; Figure 5I), IL-8 (99% increase; Figure 5J) and MCP-1 (674% increase; Figure 5K) 96 h after miRNA-145 transfection were unequivocally higher.

**Figure 5.**
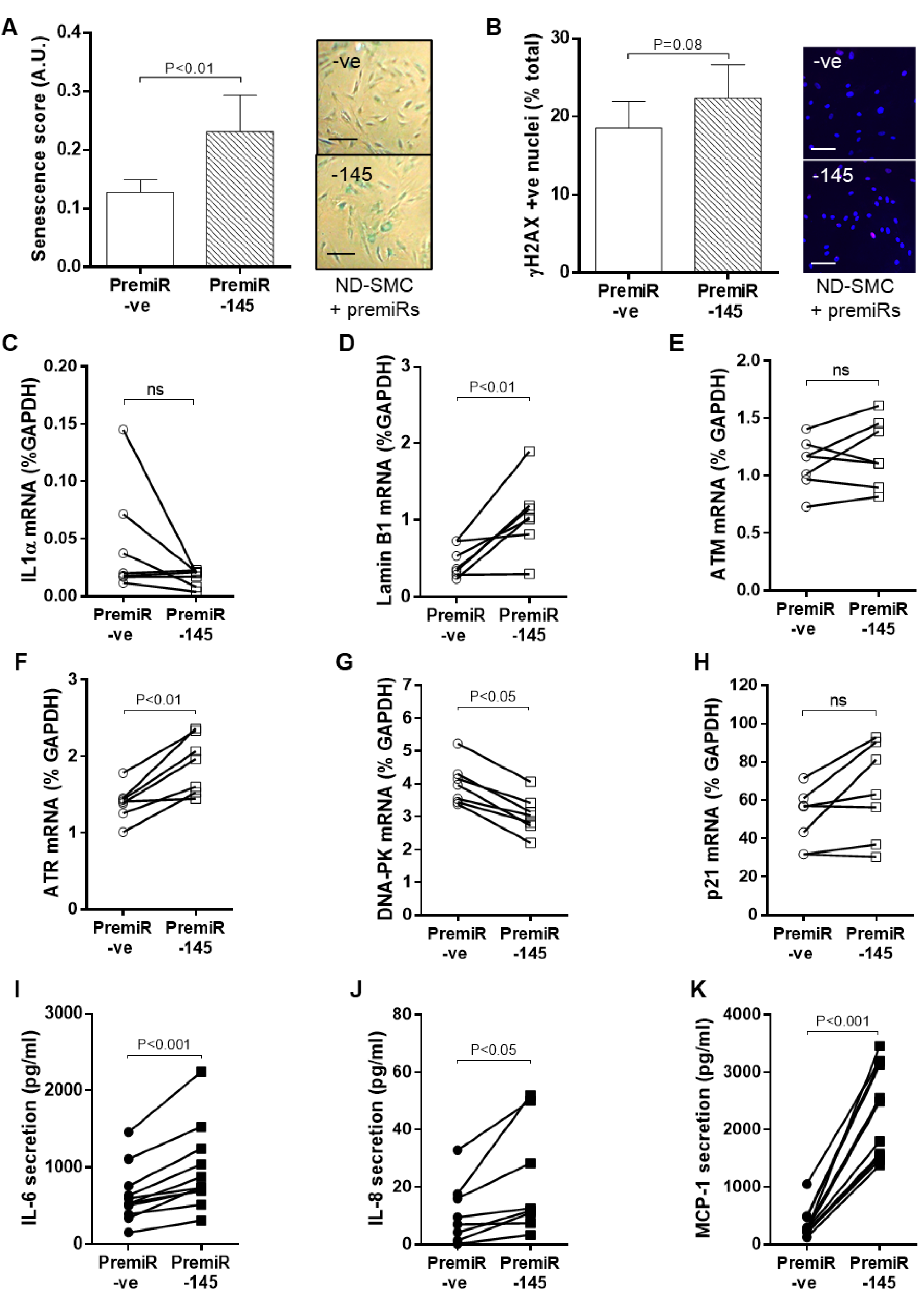
Effect of miRNA-145 overexpression on SMC senescence, DDR signalling and SASP. ND-SMC were transfected with premiRNA-145 or premiRNA -ve. **A.** After 4 days in full growth media, cells were fixed and stained for SA-β-galactosidase (scale bar = 200μM, n=6). **B.** IL-1α and **C.** LMNB1 expression were measured by RT-PCR after 72 h (n=7). **D.** Cells were treated as (A) and stained for γH2AX (pink foci, scale bar = 100μM, n=6). **E.** Expression levels of ATM, **F.** ATR, **G.** DNA-PK and **H.** p21 were quantified by RT-PCR 72 h after miRNA-145 transfection (n=7). **I.** Protein secretion of IL-6, **J.** IL-8 and **K.** MCP-1 was measured by ELISA after 96 h (n=11).

### Chronic signalling pathway activation in miRNA-145-overexpressing SMC

Persistently elevated levels of miRNA-145 in T2DM-SMC may drive chronic stress signalling to reduce proliferation. To this end we explored activation of four intracellular signalling pathways, p38MAPK, ERK, PI3K/Akt and NF-κB. Cell lysates were prepared from ND-SMC transfected 96 h earlier with premiRNA-145 or premiRNA –ve (Figure 6A). Phosphorylation of p38α was consistently elevated in miRNA-145 overexpressing cells, independent of altered p38 protein expression (Figure 6B). Increased ERK phosphorylation was accompanied by increased total protein levels (Figure 6C). miRNA-145 overexpression did not lead to changes in Akt phosphorylation or expression (Figure 6D) whereas IκB levels increased (Figure 6E), indicating reduced NF-κB pathway signalling. Finally, p21 protein expression was increased in response to miRNA-145 overexpression (Figure 6F).

**Figure 6.**
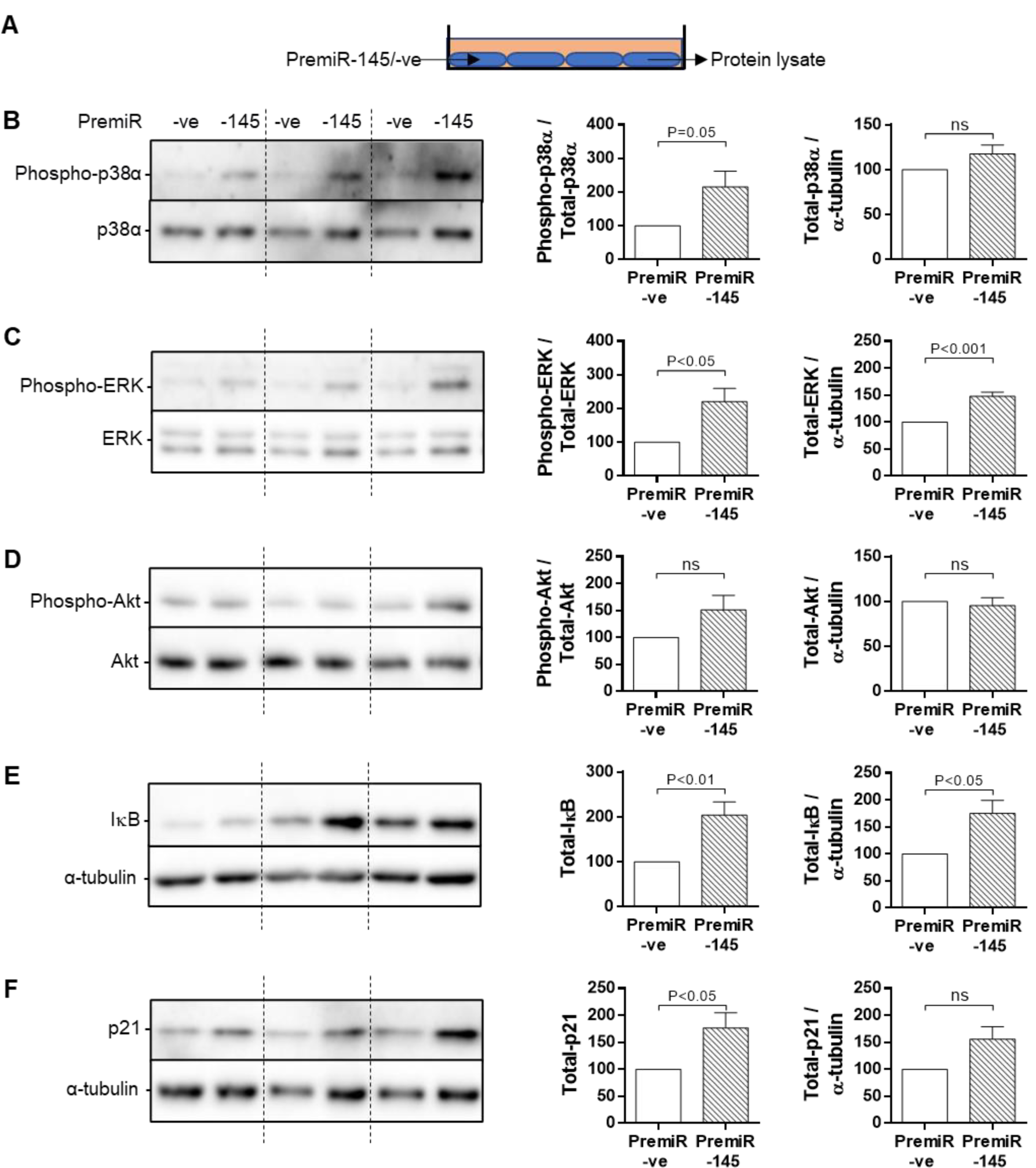
miRNA-145 overexpression induces chronic activation of intracellular signalling. **A.** Cells were transfected with premiRNA-145 or premiRNA -ve for 6 h, placed into 0.4% FCS for 96 h and protein lysates prepared. These were immunoblotted for **B.** p38, **C.** ERK, **D.** Akt (all expression and phosphorylation), **E.** IкB, **F.** p21 expression. Phosphorylation of proteins was measured against expression of the same protein, whereas changes in expression only were measured against α-tubulin as a loading control (n=6-9).

### Bystander effect of conditioned medium from miRNA-145 overexpressing cells

Senescent cells can induce senescence and DDR in neighbouring cells through paracrine mechanisms known as a “bystander effect”. ND-SMC were transfected with premiRNA –ve (control) or premiRNA-145 (overexpression) for 96 h, after which conditioned medium (CM) was collected and applied to previously serum-starved (72 h), naïve ND-SMC. After 96 h, lysates were prepared from the CM-stimulated cells and immunoblotted (Figure 7A). Again, increased phosphorylation of p38 was observed without increased total protein (Figure 7B). ERK phosphorylation was variable and inconsistent (Figure 7C). In contrast to the original miRNA-145 overexpressing cells, an increase in Akt phosphorylation was observed in the CM-treated cells in the absence of changes in total Akt (Figure 7D). IκB was unaffected (Figure 7E) and p21 protein was increased in the CM-stimulated cells, analogous to miRNA-145 transfected cells (Figure 7F).

**Figure 7.**
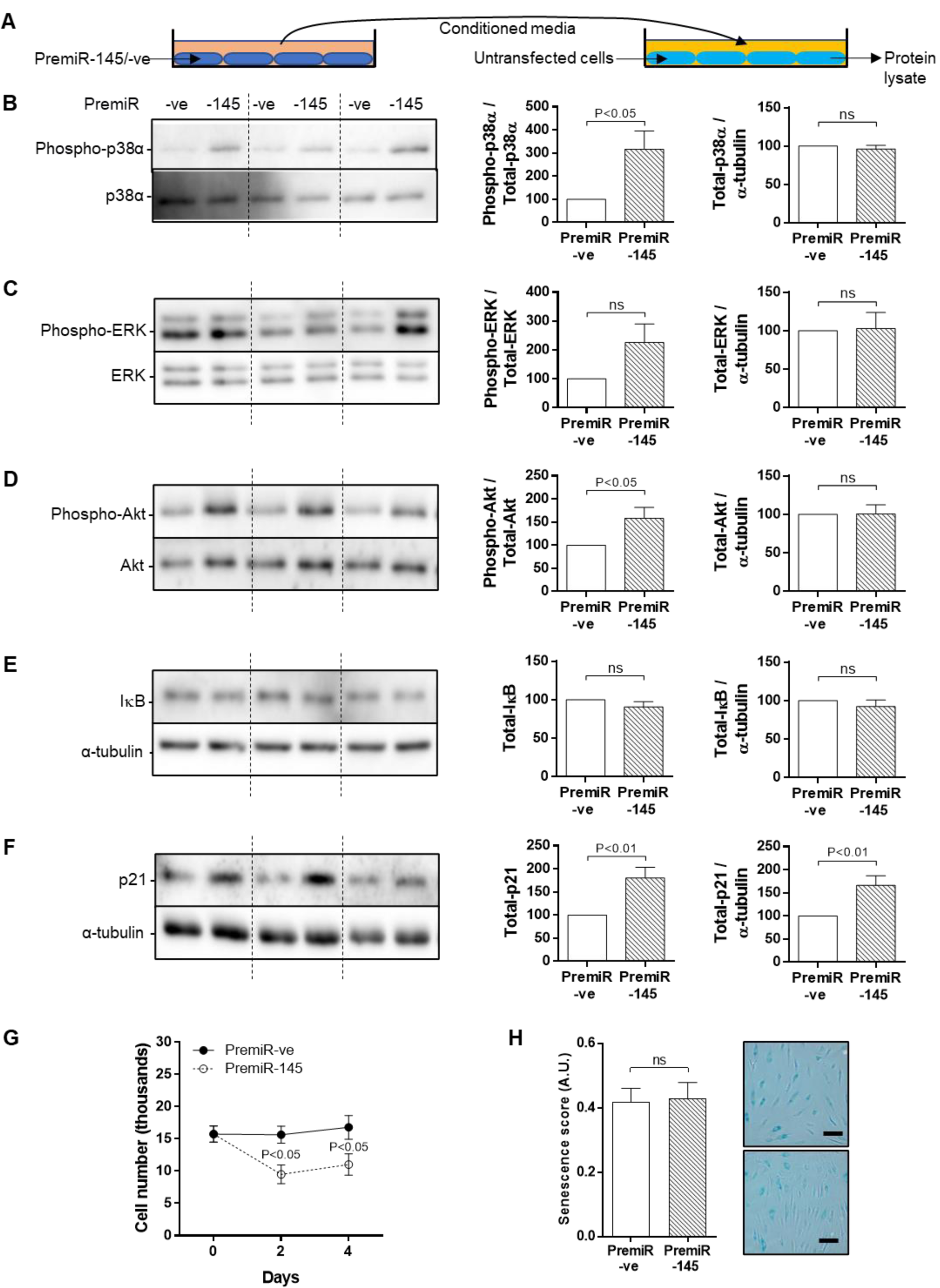
Conditioned medium from miRNA-145 overexpressing SMC induces a bystander effect in naïve cells. **A.** Cells were serum-starved for 72 h prior to exposure to conditioned (0.4% FCS) medium harvested from cells transfected with premiRNA-145 or premiRNA -ve for 96 h. Protein lysates were immunoblotted for **B.** p38, **C.** ERK, **D.** Akt (all expression and phosphorylation), **E.** IкB, **F.** p21 expression. Phosphorylation of proteins was measured against expression of the same protein, whereas changes in expression only were measured against α-tubulin as a loading control (n=6-9). **G.** Cells were serum starved for 72hrs prior to exposure to conditioned (0.4% FCS) medium harvested from cells transfected with premiRNA-145 or premiRNA -ve for 96 h. Cell counts were performed at 48 and 96h post exposure to determine cell number or (H) were fixed and stained for presence of SA-β-galactosidase staining (scale bar = 100μm; n=6-9).

Proliferation assays were performed over 4 days by exposing serum-starved naïve cells to CM from premiRNA –ve or premiRNA-145 transfected cells. Whilst control cells remained viable they did not proliferate, however significant cell loss was observed in SMC exposed to CM collected from miRNA-145 overexpressing cells (Figure 7G). Both populations exhibited substantial, yet comparable degrees of senescence (Figure 7H).

## Discussion

We previously discovered that saphenous vein SMC (SV-SMC) cultured from patients with T2DM display a persistent enlarged, flattened morphology with impaired proliferation (6) and that this was driven by an epigenetic mechanism involving elevated expression of miRNA-145 (8). Indeed, in the latter study, we demonstrated that miRNA-145 is a critical regulator of this phenotypic switching. Importantly, miRNA-145 has been identified as a tumour suppressor miRNA, confirming an association with DNA damage in a variety of cancer cell-types (17,18).

The average age of the ND and T2DM patient cohorts in that study was similar (65.2 vs 63.0 years respectively) (8); it was therefore unsurprising to reveal evidence of cellular aging in both patient populations. However, T2DM-SMC cells were unequivocally distinct from ND cells, lending weight to the idea that additional mechanisms conferring accelerated aging and senescence in T2DM may be responsible. In support of this hypothesis, we recently demonstrated in cultured human SMC of both aortic and venous origin, that miRNA-145 expression levels did not correlate with chronological age but were elevated in abdominal aortic aneurysm SMC; a disorder associated with accelerated vascular aging (19).

We proceeded to explore components of the DDR and confirmed that T2DM-SMC exhibited persistent senescence and DNA damage evidenced by aberrant cell nuclei, increased phosphorylation of H2AX and increased gene expression of ATM, ATR and p21. Whilst higher mRNA expression levels of IL-8 suggested a developing SASP, this was not corroborated by IL-6 or MCP-1 mRNA expression, or increased protein secretion suggesting that T2DM-SMC had become senescent without progressing to the secretory phenotype (20,21). It was therefore logical to explore whether inflicting DNA damage *per se* in ND-SMC could drive the acquisition of a native T2DM-SMC phenotype. In doing so, we confirmed comparable changes in proliferation, senescence, apical kinase and miRNA-145 expression. Taken together, our data indicate that T2DM-SMC exhibit enhanced DNA damage and senescence and that stimulating DNA damage in ND-SMC creates these features that are characteristic of T2DM-SMC. Our findings concur with a very recent study in which DNA damage and senescence in venous SMC was higher in T2DM versus ND patients who underwent lower limb amputation (22). The authors revealed that the observed enhanced DNA damage in the vasculature of T2DM patients plays an important role in venous SMC calcification.

Pharmacological and siRNA-mediated modulation of the apical kinases revealed that ATR inhibition led to reduced SV-SMC proliferation. This unanticipated reduction in cell number is not without precedent and has previously been documented in a variety of cancer cell lines; reportedly by inducing apoptosis (23). The reduction in cell number that we observed in proliferation assays may be reflective of enhanced apoptosis rather than impaired proliferation, although this was not evaluated. Nevertheless, neither pharmacological nor gene-silencing approaches to apical kinase inhibition modulated miRNA-145 expression, suggesting that the aberrant miRNA-145 expression was not directly driven by apical kinase activity and its effect on reducing cell number is mediated via an alternate mechanism.

Whilst accumulating evidence supports the idea that miRNAs may be novel players in the DDR (9), a functional link between miRNA-145, SMC senescence and T2DM has not been established until now. Overexpression of miRNA-145 increased β-galactosidase to levels comparable with those in native T2DM cells whilst expression patterns of IL-1α were not altered and LMNB1 was significantly increased, contrary to the reduction observed in native T2DM cells. However, published reports are at variance, for example knockdown of LMNB1 in mouse embryonic fibroblasts reduced miRNA-145 levels (24), whilst overexpression of miRNA-145 led to a reduction in LMNB1 in mesothelioma cells (25) but not in rat cardiac myocytes (26). Thus, the increased LMNB1 expression that we observed in miRNA-145 overexpressing SV-SMC may not be involved in induction of senescence.

With respect to evidence for SASP, miRNA-145 overexpressing cells secreted significantly higher levels of IL-6, IL-8 and MCP-1, supporting the idea that miRNA-145 may propagate senescence in SV-SMC via autocrine or paracrine mechanisms. In contrast, a role for miRNA-145 in the *initiation* of DNA damage was not confirmed in terms of changes in γH2AX, ATM and p21, although elevation of ATR mRNA was significant, akin to native T2DM-SMC. Taken together, these data suggest that miRNA-145 lies downstream of the initiation of DNA damage and plays a key role in provoking/reinforcing senescence.

Whilst the DDR pathway is rapid, development of SASP is a delayed and often prolonged event over days or weeks (27,28), inferring that canonical DDR signalling is insufficient to drive SASP and that additional molecular mechanisms are necessary. Notably, the p38 MAPK pathway has been shown to be a signalling target of a variety of cellular stresses – oxidative, metabolic, DNA damage and mechanical damage (27). Importantly, p38 sustains SASP in cancer cells and tumorigenesis (29) and has been described as a marker of vascular SMC senescence (21).

Our studies revealed chronic activation of p38α in miRNA-145 overexpressing cells. A complex, bi-directional relationship between miRNA-145 and p38 appears to exist. For example, p38 phosphorylation was suppressed by miRNA-145 in placental tissue (30), yet in a different study an increase in p38 phosphorylation was noted on overexpression *and* knockdown of miRNA-145 in chondrocytes (31). A p38 response element has been identified within the promoter of miRNA-145 (32,33), and p38 increased the processing of pri-miRNA-145 to mature miRNA-145 (34).

We also observed increased ERK1/2 phosphorylation that was accompanied by increased ERK protein expression. The relationship between miRNA-145 and ERK1/2 in general appears to be inverse; indeed ERK1/2 inhibited miRNA-145 expression in human aortic SMC (35) and miRNA-145 inhibition reportedly enhanced ERK1/2 and Akt phosphorylation in cancer cells (36). Conversely, we observed a positive relationship between miRNA-145 and ERK1/2 which might be indicative of a negative feedback mechanism. Similarly, the observed increase in the anti-inflammatory protein IкB may be a compensatory mechanism by the cell to ameliorate stress signalling. Taken together, these observations provide evidence of chronic stress in miRNA-145 overexpressing SV-SMC.

p21 is a key downstream effector of p53 in the DDR pathway, halting the cell cycle to allow DNA damage to be repaired (37). We observed significantly elevated p21 expression in native T2DM cells which was mimicked by inducing DNA damage or over-expressing miRNA-145. This is in keeping with reduced proliferation in T2DM and miRNA-145 over-expressing cells, and concurs with previous literature where miRNA-145 and p53 have a complementary positive feedback relationship that increases p21 expression (38).

Factors released from senescent cells can inflict an autocrine or paracrine response in neighbouring cells, known as a bystander effect (39), that can further reinforce DNA damage and senescence. We showed that activation of p38 and expression of p21 were analogous to that observed in “donor” miRNA-145 overexpressing cells, suggesting that these signalling events are important for both induction and propagation of DNA damage and cellular senescence.

In summary, emerging evidence supports a role for epigenetics in VSMC senescence and aging and in particular, proposed functions for miRNAs [reviewed in (21)]. Importantly, we have identified a novel mechanism linking aberrant miRNA-145 expression to SV-SMC senescence in the setting of macrovascular complications of T2DM. Figure 8 provides an illustrative summary of our findings. Succinctly, miRNA-145 drives increased senescence, reduced cell proliferation and activation of chronic stress signalling in response to DNA damage although its potential to instigate DNA damage may be secondary to these important aspects.

**Figure 8.**
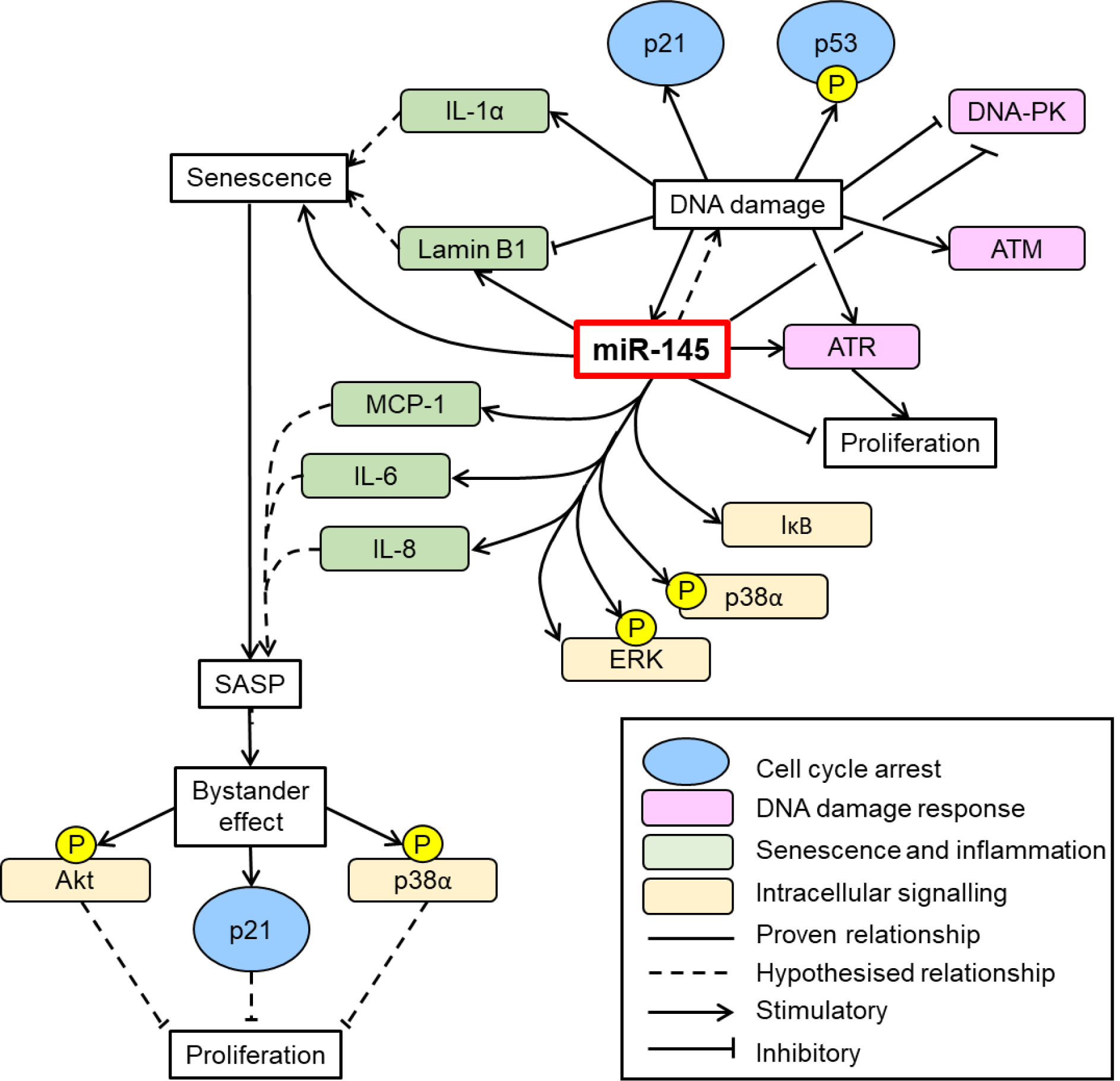
Summary of the central role for miRNA-145 in influencing SV-SMC DNA damage, proliferation and senescence. DNA damage upregulates expression of miRNA-145 in addition to inducing cell cycle arrest (p21 and p53 phosphorylation), upregulating apical kinase expression (ATR, ATM, DNA-PK) and cellular senescence (IL-1α, Lamin B1 and SA β-gal. miRNA-145 reinforces DDR by also upregulating ATR expression, triggering intracellular kinase cascades (IкB, p38α, ERK) and inducing components of SASP (MCP-1, IL-6, IL-8). This induction of senescence and SASP inflicts a paracrine bystander effect on neighbouring cells through chronic activation of Akt and p38α, which can block cellular proliferation and perpetuate the T2DM-SMC phenotype.

## Experimental procedures

### Smooth muscle cell (SMC) culture

SMC were cultured as previously described (40) from explants of SV obtained from non-diabetic patients (ND-SMC), or patients diagnosed with type 2 diabetes (T2DM-SMC) receiving oral therapy either alone or with the addition of insulin. Patients had similar age and gender profiles and were undergoing elective CABG surgery at the Leeds General Infirmary or Nuffield Health Leeds hospital. Samples were collected with local ethical committee approval and informed, written patient consent. The study conformed to the principles outlined in the Declaration of Helsinki. SMC were cultured in DMEM containing 10% bovine FCS, 25 mmol/L glucose, 1% penicillin, streptomycin, fungizone and 2 mmol/L glutamine (full growth medium; FGM) at 37°C in 5% CO2 in air. All experiments were performed on cells between passages 3 to 6.

### Senescence associated β-galactosidase staining

SMC were seeded at a density of 75000/well in 6 well plates and cultured in FGM for 48 h prior to fixation and staining using a commercial β-galactosidase assay (Cell Signaling Technologies #9860) as described previously (41). Senescent cells were identified by formation of a blue precipitate at pH 6 and senescence scores determined by counting the proportion of senescent cells contained within 10 low power (x 40) microscopic fields of view.

### Quantitative real-time RT-PCR

Cellular RNA was extracted and cDNA prepared as previously described (42). Real-time RT-PCR was performed in triplicate using an Applied Biosystems 7500 Real-time PCR system and Taqman Gene Expression Assays (Thermo-Fisher Scientific). Data are presented as percentage of endogenous control (GAPDH) expression using the formula 2^-ΔCT^ x 100.

### Apoptosis Assay

SMC were plated in FGM at a density of 3000/well in 96 well plates and incubated overnight. Apoptosis assays were performed as previously described (41). Briefly, cells were treated with 5 μmol/L NucView™ 488 caspase-3 substrate (Biotium). Images were obtained in phase contrast and fluorescence mode using a x10 objective and an IncuCyte FLR time-lapse fluorescence microscope (Essen Bioscience). After 24 h, cells were stained with 1 μmol/L Vybrant DyeCycle Green® (Molecular Probes, Invitrogen) and an apoptosis index calculated using an inbuilt algorithm.

### Immunocytochemistry

SMC cultured on glass coverslips were fixed in 4% PFA prior to immunostaining with phospho-histone H2AX (γH2AX) antibody (Cell Signaling Technologies #2577) as described previously (19). Coverslips were mounted using Prolong Gold antifade reagent containing DAPI (Thermo-Fisher Scientific) and imaged on a Zeiss 700 confocal microscope. Nuclear morphology was examined in DAPI-stained cells at x400 magnification and the proportion of cell nuclei displaying DNA damage (blebbing or apoptotic nuclei) was quantified as the number of γH2AX-positive nuclei (pink) relative to the total number of regular, ovoid nuclei (blue).

### Enzyme linked immunosorbent assays (ELISA)

Cell culture supernatants were collected from SMC, snap frozen and stored at -80°C. Secretion of IL-6, IL-8 and MCP-1 was determined by ELISA (Quantikine, R&D Systems).

### Cell proliferation

Cells were seeded in 24-well plates at a density of 10000 cells/well and triplicate counts performed over a 7-day period using a haemocytometer and trypan blue exclusion, as previously (6).

### Immunoblotting

Whole cell homogenates were prepared and immunoblotted as described previously (42) using antibodies to p-p38 (#9215), total p38 (#9228), p-ERK (#9106), total ERK (#4695), p-Akt (#4691), total Akt (#2920), IκB (#9242), p21 (#2947), p-p53 (ser15) (#9284) (all Cell Signaling Technologies), with α-tubulin (ab8226; Abcam) as a loading control.

### Quantification of miRNA-145 levels

Total RNA samples were analysed by preparing specific RT reactions for U6 and miRNA-145 using Taqman MicroRNA Assays (Applied Biosystems). Real-time RT-PCR was performed in triplicate and data presented as percentage endogenous control (U6) expression using the formula 2^-ΔCT^ x 100.

### Inhibition of DDR apical kinases

Cells were seeded in FGM at 10000 cells/well in 24 well plates, cultured overnight and then serum-deprived for 72 h prior to re-addition of FGM plus inhibitors of ATM (KU55933, 10 μM), ATR (AZ20, 10 μM), DNA-PK (NU7026, 3 μM) (all Generon) or DMSO vehicle.

### siRNA knockdown

SMC were transfected with siRNA to ATM, ATR, DNA-PK or scrambled control (20 nM, ONTARGETplus siRNA, Dharmacon) using Lipofectamine 2000 reagent (Thermo Fisher Scientific), according to the manufacturer’s instructions.

### miRNA-145 overexpression

Overexpression of miRNA-145 was achieved by transfecting SMC with 15 nM premiRNA-145 or negative control (premiRNA –ve) as described previously (8).

### Statistical analysis

Statistical analysis was performed using GraphPad Prism 7 software. Data are presented as mean±SEM with *n* representing the number of experiments performed on cells from different patients. Data were tested for normality prior to log transformation and analysis by paired or unpaired t-test, or one way ANOVA with Dunnett’s post-hoc test, as appropriate. P<0.05 was considered significant.

### Data availability

All data are contained within the article and supporting information. Data are available upon request.

## Footnotes

This article contains supporting information

## Funding and additional information

This work was supported by a Diabetes UK project grant (15/0005143) awarded to K.E. Porter and N.A. Turner.

## Conflict of interest

The authors declare that they have no conflicts of interest with the content of this article

## Abbreviations

The abbreviations used are:

ATM: ataxia-telangiectasia mutated
ATR: ataxia-telangiectasia mutated and Rad3 related;
CABG: coronary artery bypass grafting
CHD: coronary heart disease
CM: conditioned medium
DDR: DNA damage response
DNA-PK: DNA dependent protein kinase catalytic subunit
FGM: full growth medium
ERK: extracellular signal-regulated kinase
GAPDH: glyceraldehyde 3-phosphate dehydrogenase
IL: interleukin
LMNB1: lamin B1
MAPK: mitogen-activated protein kinase
MCP-1: monocyte chemoattractant protein-1
miRNA: microRNA
ND: non-diabetic
SASP: senescence-associated secretory phenotype
SMC: smooth muscle cell
SV: saphenous vein
T2DM: type 2 diabetes mellitus;

## References

1. Booth, G. L., Kapral, M. K., Fung, K., and Tu, J. V. (2006) Relation between age and cardiovascular disease in men and women with diabetes compared with non-diabetic people: a population-based retrospective cohort study. Lancet 368, 29–36

2. Katsuumi, G., Shimizu, I., Yoshida, Y., and Minamino, T. (2018) Vascular Senescence in Cardiovascular and Metabolic Diseases. Front. Cardiovasc. Med 5, 18

3. Hakala, T., Pitkanen, O., Halonen, P., Mustonen, J., Turpeinen, A., and Hippelainen, M. (2005) Early and late outcome after coronary artery bypass surgery in diabetic patients. Scand. Cardiovasc. J 39, 177–181

4. Kubal, C., Srinivasan, A. K., Grayson, A. D., Fabri, B. M., and Chalmers, J. A. (2005) Effect of risk-adjusted diabetes on mortality and morbidity after coronary artery bypass surgery. Ann. Thorac. Surg 79, 1570–1576

5. Owens, C. D. (2010) Adaptive changes in autogenous vein grafts for arterial reconstruction: clinical implications. J. Vasc. Surg 51, 736–746

6. Madi, H. A., Riches, K., Warburton, P., O’Regan, D. J., Turner, N. A., and Porter, K. E. (2009) Inherent differences in morphology, proliferation, and migration in saphenous vein smooth muscle cells cultured from nondiabetic and Type 2 diabetic patients. Am. J. Physiol Cell Physiol 297, C1307–C1317

7. Riches, K., Warburton, P., O’Regan, D. J., Turner, N. A., and Porter, K. E. (2014) Type 2 diabetes impairs venous, but not arterial smooth muscle cell function: Possible role of differential RhoA activity. Cardiovasc. Revasc. Med 15, 141–148

8. Riches, K., Alshanwani, A. R., Warburton, P., O’Regan, D. J., Ball, S. G., Wood, I. C., Turner, N. A., and Porter, K. E. (2014) Elevated expression levels of miR-143/5 in saphenous vein smooth muscle cells from patients with Type 2 diabetes drive persistent changes in phenotype and function. J. Mol. Cell Cardiol 74, 240–250

9. Hu, H., and Gatti, R. A. (2011) MicroRNAs: new players in the DNA damage response. J. Mol. Cell Biol 3, 151–158

10. Ugalde, A. P., Ramsay, A. J., de la Rosa, J., Varela, I., Marino, G., Cadinanos, J., Lu, J., Freije, J. M., and Lopez-Otin, C. (2011) Aging and chronic DNA damage response activate a regulatory pathway involving miR-29 and p53. EMBO J 30, 2219–2232

11. Iaconetti, C., De, R. S., Polimeni, A., Sorrentino, S., Gareri, C., Carino, A., Sabatino, J., Colangelo, M., Curcio, A., and Indolfi, C. (2015) Down-regulation of miR-23b induces phenotypic switching of vascular smooth muscle cells in vitro and in vivo. Cardiovasc. Res 107, 522–533

12. Santulli, G., Wronska, A., Uryu, K., Diacovo, T. G., Gao, M., Marx, S. O., Kitajewski, J., Chilton, J. M., Akat, K. M., Tuschl, T., Marks, A. R., and Totary-Jain, H. (2014) A selective microRNA-based strategy inhibits restenosis while preserving endothelial function. J. Clin. Invest 124, 4102–4114

13. Torella, D., Iaconetti, C., Catalucci, D., Ellison, G. M., Leone, A., Waring, C. D., Bochicchio, A., Vicinanza, C., Aquila, I., Curcio, A., Condorelli, G., and Indolfi, C. (2011) MicroRNA-133 controls vascular smooth muscle cell phenotypic switch in vitro and vascular remodeling in vivo. Circ. Res 109, 880–893

14. Wang, Y., and Taniguchi, T. (2013) MicroRNAs and DNA damage response: implications for cancer therapy. Cell Cycle 12, 32–42

15. Bennett, M. R., and Clarke, M. C. (2017) Killing the old: cell senescence in atherosclerosis. Nat. Rev. Cardiol 14, 132

16. Freund, A., Laberge, R. M., Demaria, M., and Campisi, J. (2012) Lamin B1 loss is a senescence-associated biomarker. Mol. Biol. Cell 23, 2066–2075

17. Pashaei, E., Guzel, E., Ozgurses, M. E., Demirel, G., Aydin, N., and Ozen, M. (2016) A Meta-Analysis: Identification of Common Mir-145 Target Genes that have Similar Behavior in Different GEO Datasets. PloS one 11, e0161491

18. Suzuki, H. I., Yamagata, K., Sugimoto, K., Iwamoto, T., Kato, S., and Miyazono, K. (2009) Modulation of microRNA processing by p53. Nature 460, 529–533

19. Riches, K., Clark, E., Helliwell, R. J., Angelini, T. G., Hemmings, K. E., Bailey, M. A., Bridge, K. I., Scott, D. J. A., and Porter, K. E. (2018) Progressive Development of Aberrant Smooth Muscle Cell Phenotype in Abdominal Aortic Aneurysm Disease. J. Vasc. Res 55, 35–46

20. Noren Hooten, N., and Evans, M. K. (2017) Techniques to Induce and Quantify Cellular Senescence. J. Vis. Exp 123, 55533

21. Chi, C., Li, D. J., Jiang, Y. J., Tong, J., Fu, H., Wu, Y. H., and Shen, F. M. (2019) Vascular smooth muscle cell senescence and age-related diseases: State of the art. Biochim. Biophys. Acta. Mol. Basis. Dis 1865, 1810–1821

22. Bartoli-Leonard, F., Wilkinson, F. L., Schiro, A., Inglott, F. S., Alexander, M. Y., and Weston, R. (2020) Loss of SIRT1 in diabetes accelerates DNA damage induced vascular calcification. Cardiovasc. Res doi: 10.1093/cvr/cvaa134

23. Job, A., Schmitt, L. M., von Wenserski, L., Lankat-Buttgereit, B., Gress, T. M., Buchholz, M., and Gallmeier, E. (2018) Inactivation of PRIM1 Function Sensitizes Cancer Cells to ATR and CHK1 Inhibitors. Neoplasia (New York, N.Y.) 20, 1135–1143

24. Malhas, A., Saunders, N. J., and Vaux, D. J. (2010) The nuclear envelope can control gene expression and cell cycle progression via miRNA regulation. Cell Cycle 9, 531–539

25. Cioce, M., Ganci, F., Canu, V., Sacconi, A., Mori, F., Canino, C., Korita, E., Casini, B., Alessandrini, G., Cambria, A., Carosi, M. A., Blandino, R., Panebianco, V., Facciolo, F., Visca, P., Volinia, S., Muti, P., Strano, S., Croce, C. M., Pass, H. I., and Blandino, G. (2014) Protumorigenic effects of mir-145 loss in malignant pleural mesothelioma. Oncogene 33, 5319–5331

26. Li, R., Yan, G., Zhang, Q., Jiang, Y., Sun, H., Hu, Y., Sun, J., and Xu, B. (2013) miR-145 inhibits isoproterenol-induced cardiomyocyte hypertrophy by targeting the expression and localization of GATA6. FEBS Lett 587, 1754–1761

27. Freund, A., Patil, C. K., and Campisi, J. (2011) p38MAPK is a novel DNA damage response-independent regulator of the senescence-associated secretory phenotype. EMBO J 30, 1536–1548

28. Malaquin, N., Carrier-Leclerc, A., Dessureault, M., and Rodier, F. (2015) DDR-mediated crosstalk between DNA-damaged cells and their microenvironment. Front. Genet 6, 94

29. Alspach, E., Flanagan, K. C., Luo, X., Ruhland, M. K., Huang, H., Pazolli, E., Donlin, M. J., Marsh, T., Piwnica-Worms, D., Monahan, J., Novack, D. V., McAllister, S. S., and Stewart, S. A. (2014) p38MAPK plays a crucial role in stromal-mediated tumorigenesis. Cancer Discov 4, 716–729

30. Farrokhnia, F., Aplin, J. D., Westwood, M., and Forbes, K. (2014) MicroRNA regulation of mitogenic signaling networks in the human placenta. J. Biol. Chem 289, 30404–30416

31. Hu, G., Zhao, X., Wang, C., Geng, Y., Zhao, J., Xu, J., Zuo, B., Zhao, C., Wang, C., and Zhang, X. (2017) MicroRNA-145 attenuates TNF-alpha-driven cartilage matrix degradation in osteoarthritis via direct suppression of MKK4. Cell Death Dis 8, e3140

32. Long, X., and Miano, J. M. (2011) Transforming growth factor-beta1 (TGF-□1) utilizes distinct pathways for the transcriptional activation of microRNA 143/145 in human coronary artery smooth muscle cells. J. Biol. Chem 286, 30119–30129

33. O’Leary, L., Sevinc, K., Papazoglou, I. M., Tildy, B., Detillieux, K., Halayko, A. J., Chung, K. F., and Perry, M. M. (2016) Airway smooth muscle inflammation is regulated by microRNA-145 in COPD. FEBS Lett 590, 1324–1334

34. Hong, S., Noh, H., Chen, H., Padia, R., Pan, Z. K., Su, S. B., Jing, Q., Ding, H. F., and Huang, S. (2013) Signaling by p38 MAPK stimulates nuclear localization of the microprocessor component p68 for processing of selected primary microRNAs. Sci. Signal 6, ra16

35. Hu, B., Wu, Z., Jin, H., Hashimoto, N., Liu, T., and Phan, S. H. (2004) CCAAT/enhancer-binding protein β isoforms and the regulation of β-smooth muscle actin gene expression by IL-1?. J. Immunol 173, 4661–4668

36. Yin, Y., Yan, Z. P., Lu, N. N., Xu, Q., He, J., Qian, X., Yu, J., Guan, X., Jiang, B. H., and Liu, L. Z. (2013) Downregulation of miR-145 associated with cancer progression and VEGF transcriptional activation by targeting N-RAS and IRS1. Biochim. Biophys. Acta 1829, 239–247

37. Bunz, F., Dutriaux, A., Lengauer, C., Waldman, T., Zhou, S., Brown, J. P., Sedivy, J. M., Kinzler, K. W., and Vogelstein, B. (1998) Requirement for p53 and p21 to sustain G2 arrest after DNA damage. Science 282, 1497–1501

38. Spizzo, R., Nicoloso, M. S., Lupini, L., Lu, Y., Fogarty, J., Rossi, S., Zagatti, B., Fabbri, M., Veronese, A., Liu, X., Davuluri, R., Croce, C. M., Mills, G., Negrini, M., and Calin, G. A. (2010) miR-145 participates with TP53 in a death-promoting regulatory loop and targets estrogen receptor-alpha in human breast cancer cells. Cell Death Diff 17, 246–254

39. Hubackova, S., Krejcikova, K., Bartek, J., and Hodny, Z. (2012) IL1- and TGFbeta-Nox4 signaling, oxidative stress and DNA damage response are shared features of replicative, oncogene-induced, and drug-induced paracrine ‘bystander senescence’. Aging (Albany. NY) 4, 932–951

40. Porter, K. E., Naik, J., Turner, N. A., Dickinson, T., Thompson, M. M., and London, N. J. (2002) Simvastatin inhibits human saphenous vein neointima formation via inhibition of smooth muscle cell proliferation and migration. J. Vasc. Surg 36, 150–157

41. Riches, K., Angelini, T. G., Mudhar, G. S., Kaye, J., Clark, E., Bailey, M. A., Sohrabi, S., Korossis, S., Walker, P. G., Scott, D. J., and Porter, K. E. (2013) Exploring smooth muscle phenotype and function in a bioreactor model of abdominal aortic aneurysm. J. Transl. Med 11, 208

42. Turner, N. A., Mughal, R. S., Warburton, P., O’Regan, D. J., Ball, S. G., and Porter, K. E. (2007) Mechanism of TNFα-induced IL-1α, IL-1β and IL-6 expression in human cardiac fibroblasts: Effects of statins and thiazolidinediones. Cardiovasc. Res 76, 81–90

